# Cell-type specific D1 dopamine receptor modulation of projection neurons and interneurons in the prefrontal cortex

**DOI:** 10.1101/370536

**Authors:** Paul G. Anastasiades, Christina Boada, Adam G. Carter

## Abstract

Dopamine modulation in the prefrontal cortex (PFC) mediates diverse effects on neuronal physiology and function, but the expression of dopamine receptors at sub-populations of pyramidal neurons and interneurons remains unresolved. Here, we examine D1 receptor expression and modulation at specific cell types and layers in the mouse prelimbic PFC. We first show that D1 receptors are enriched in pyramidal neurons in both layers 5 and 6, and that these cells project intra-telencephalically, rather than to sub-cortical structures. We then find that D1 receptors are also present in interneurons, and enriched in VIP+ interneurons that co-expresses calretinin, but absent from PV+ and SOM+ interneurons. Finally, we determine that D1 receptors strongly and selectively enhance action potential firing in only a subset of these cortico-cortical neurons and VIP+ interneurons. Our findings define several novel sub-populations of D1+ neurons, highlighting how modulation via D1 receptors can influence both excitatory and disinhibitory micro-circuits in the PFC.

## INTRODUCTION

Dopamine modulation in the prefrontal cortex (PFC) plays a key role in cognitive, motivational and emotional behavior (Fuster JM 2000; Miller EK and JD Cohen 2001). Dopamine can signal through multiple receptors, which are subdivided into D1-like (D1 and D5) or D2-like (D2, D3 and D4) (Tritsch NX and BL Sabatini 2012). While all these subtypes (D1-5) are present in the PFC (Weiner DM et al. 1991; Mrzljak L et al. 1996; Oda S et al. 2010; Clarkson RL et al. 2017), D1 receptors (D1-Rs) are the most abundant (Lidow MS et al. 1991; Gaspar P et al. 1995; Santana N et al. 2009). The importance of D1-R signaling is highlighted by its requirement for PFC-dependent behaviors (Sawaguchi T and PS Goldman-Rakic 1994; Williams GV and PS Goldman-Rakic 1995; Seamans JK et al. 1998), and its disruption in many neuropsychiatric disorders (Grace AA 2016). Determining which cell types express D1-Rs is therefore essential for understanding how dopamine modulates the PFC in health and disease.

The ability of the PFC to mediate executive control ultimately depends on the diverse long-range projections it sends to other brain regions (Miller EK and JD Cohen 2001; Gabbott PL et al. 2005). For example, intra-telencephalic (IT) neurons project within the cortex, including between cerebral hemispheres, and are distinct from pyramidal tract (PT) neurons that project sub-cortically (Gabbott PL *et al.* 2005; Dembrow NC et al. 2010; Anastasiades PG et al. 2018). Recent studies indicate that dopamine receptors may differentially segregate between these two broad populations of layer 5 projection neurons in the PFC (Gee S et al. 2012; Seong HJ and AG Carter 2012; Clarkson RL *et al.* 2017). Interestingly, D1-Rs are also expressed in layer 6 (L6), where they may modulate cortico-thalamic (CT) projections (Gaspar P *et al.* 1995). However, there is currently no consensus on which projection neurons primarily express D1-Rs, with reports varying significantly (Vincent SL et al. 1993; Gaspar P *et al.* 1995). Given that the activity of defined projection neurons can have distinct effects on behavior (Land BB et al. 2014; Jenni NL et al. 2017; Murugan M et al. 2017; Otis JM et al. 2017), this represents a significant gap in our understanding of how dopamine modulates PFC outputs.

Like other cortices, the PFC also contains a diverse array of GABAergic interneurons (Markram H et al. 2004; Petilla Interneuron Nomenclature G et al. 2008; Anastasiades PG and SJ Butt 2011; Rudy B et al. 2011). Interneurons are segregated into distinct subtypes based on their intrinsic electrophysiology, morphology, and immunohistochemical markers (Kubota Y and Y Kawaguchi 1994; Kawaguchi Y and Y Kubota 1996; Butt SJ et al. 2005; Gonchar Y et al. 2007; Xu X et al. 2010; Anastasiades PG et al. 2016). Moreover, dopamine receptors are thought to be expressed in several populations of interneurons (Muly EC, 3rd et al. 1998;Glausier JR et al. 2009; Santana N *et al.* 2009), where they can mediate diverse effects (Gonzalez-Islas C and JJ Hablitz 2001; Kroner S et al. 2007; Towers SK and S Hestrin 2008; Karunakaran S et al. 2016). However, as with projection neurons, there are conflicting reports on which interneuron subtypes express D1-Rs in the PFC (Le Moine C and P Gaspar 1998; Muly EC, 3rd *et al.* 1998;Paspalas CD and PS Goldman-Rakic 2005; Santana N *et al.* 2009). While neuromodulation of inhibition plays a key role in cortical circuits (Kruglikov I and B Rudy 2008; Letzkus JJ et al. 2011; Wester JC and CJ McBain 2014; Froemke RC 2015), it is currently unclear whether D1-Rs primarily regulate inhibitory or disinhibitory networks, which is critical to informing models of PFC function (Wang XJ et al. 2004; Glausier JR *et al.* 2009).

Here we examine D1-receptor expressing (D1+) neurons in the mouse prelimbic PFC, combining *ex vivo* electrophysiology, selective pharmacology, 2-photon microscopy, immunohistochemistry and retrograde anatomy in multiple transgenic mouse lines to selectively label different populations of projection neurons and interneurons. We find that D1-Rs are strongly expressed in sub-populations of IT neurons found in both L5 and L6. Surprisingly, D1-Rs are absent from parvalbumin (PV+) and somatostatin (SOM+) expressing interneurons, but are selectively enriched in a sub-population of superficial interneurons that express vasoactive intestinal peptide (VIP+). Activation of D1-Rs enhances firing in both D1+ pyramidal neurons and VIP+ interneurons, indicating that D1-Rs enhance both excitatory and disinhibitory micro-circuits in the PFC.

## MATERIALS & METHODS

### Animals

Experiments used either heterozygous D1-tdTomato mice (Ade KK et al. 2011), or heterozygous D1-tdTomato mice crossed with either homozygous GAD-Cre (Taniguchi H et al. 2011), PV-Cre (Hippenmeyer S et al. 2005), SOM-Cre (Taniguchi H *et al.* 2011), VIP-Cre (Taniguchi H *et al.* 2011) or heterozygous 5HT3a-Cre mice (Gerfen CR et al. 2013). Mice were bred on a C57BL/6J background with the exception of D1-tdTomato x VIP-Cre mice, which were mixed background. All mice were purchased from Jackson laboratories. Mice of both sexes were used, and no differences were found. All experimental procedures were approved by the University Animal Welfare Committee of New York University.

### Stereotaxic injections

Mice aged 4-7 weeks were deeply anesthetized with a mixture of ketamine (10 mg/mL) and xylazine (0.1 mg/mL) and head fixed in a stereotax (Kopf Instruments). A small craniotomy was made over the injection site, using coordinates relative to Bregma (dorsoventral, mediolateral, and rostrocaudal, respectively): prefrontal cortex (PFC) = −2.1, ±0.4, +2.2 mm; claustrum (CLA) = −3.6, −3.2, +1.6 mm (injected at 5 degrees to the upright); mediodorsal thalamus (MD) = −3.6, −0.3, −0.5 mm; ventromedial thalamus (VM) = −3.4, −2.7, −0.4 mm (injected at 30 degrees to the upright); ventral tegmental area (VTA) = −4.5, −0.5, −2.95 mm; basolateral amygdala (BLA) = −4.9, −3.2, −1.2 mm, dorsomedial striatum (STR) = −3.2, −1.1, +1.1 mm, pontine nucleus (pons) = −4.7, +0.5, - 4.0 mm. For retrograde labeling, pipettes were filled with Cholera Toxin Subunit B (CTB) conjugated to either Alexa-488 or −647 (Life Technologies). Virus varied between experiment: Cre-dependent labeling of interneurons = AAV9-CAG-FLEX-EGFP-WPRE (UPenn); labeling putative pyramidal neurons = AAV1-CaMKII-EGFP-WPRE (UPenn); axon anatomy = AAV-DJ-hSyn1-mCherry-IRES-eGFP-Syb2 (SynaptoTag, Stanford); retrograde-Cre and associated axon anatomy = AAVrg-EF1a-mCherry-IRES-Cre (Addgene) and AAV1-EF1a-DIO-eYFP-WPRE (UPenn). Borosilicate pipettes with 5-10 μm tip diameters were backfilled, and between 130-550 nL of solution was pressure injected using a Nanoject III (Drummond), with 30 s spacing between injections. The pipette was subsequently left in place for an additional 5 min, allowing time to diffuse away from the pipette tip, before being slowly retracted from the brain. Animals were returned to their cages for between 1-3 weeks before being used for recording or anatomy, or for 4-6 weeks in the case of the SynaptoTag virus injections.

### Slice preparation

Mice aged 6-8 weeks were anesthetized with a lethal dose of ketamine (25 mg/mL) and xylazine (0.25 mg/mL) and perfused intracardially with ice-cold external solution containing the following (in mM): 65 sucrose, 76 NaCl, 25 NaHCO_3_, 1.4 NaH_2_PO_4_, 25 glucose, 2.5 KCl, 7 MgCl_2_, 0.4 Na-ascorbate, and 2 Na-pyruvate (295-305 mOsm), and bubbled with 95% O_2_/5% CO_2_. Coronal slices (300 μm thick) were cut on a VS1200 vibratome (Leica) in ice-cold external solution, before being transferred to ACSF containing the following (in mM): 120 NaCl, 25 NaHCO_3_, 1.4 NaH_2_PO_4_, 21 glucose, 2.5 KCl, 2 CaCl_2_, 1 MgCl_2_, 0.4 Na-ascorbate, and 2 Na-pyruvate (295-305 mOsm), bubbled with 95% O_2_/5% CO_2_. Slices were kept for 30 min at 35°C, before being allowed to recover for 30 min at room temperature. Intrinsic properties were recorded at 30−32°C. To facilitate stable recordings of cells with very high input resistance, modulation of VIP+ interneurons was performed at room temperature. Modulation of pyramidal cells was performed at both 30−32°C and room temperature, with no differences observed across these conditions, so results were pooled for analysis.

### Electrophysiology

Whole-cell recordings were obtained from neurons across all layers of the prelimbic subdivision of PFC. Neurons were identified by infrared-differential interference contrast, as previously described (Chalifoux JR and AG Carter 2010). Neuronal identity was established by the presence or absence of tdTomato, EGFP and Alexa-conjugated CTB under fluorescent illumination. Borosilicate pipettes (2-6 MΩ) were filled with internal solution comprising (in mM): 135 K-gluconate, 7 KCl, 10 HEPES, 10 Na-phosphocreatine, 4 Mg_2_-ATP, 0.4 Na-GTP and 0.5 EGTA, 290–295 mOsm, pH 7.3, with KOH. For a subset of experiments, 30 μM Alexa Fluor 594 was included for 2-photon imaging, in which case dye was allowed to diffuse throughout the dendrites and axons for at least 20 min before imaging.

Electrophysiology recordings were made with a Multiclamp 700B amplifier (Axon Instruments), filtered at 4 kHz, and sampled at 10 kHz. Series resistance was typically <20 MΩ for pyramidal neurons and <30 MΩ for VIP+ interneurons. Current-clamp recordings were performed in the presence of the synaptic blockers CPP (10 μM), NBQX (10 μM) and Gabazine (10 μM). Dopamine receptor pharmacology was performed using wash-in of the selective D1-type dopamine receptor agonist SKF-81297 (10 μM) and the selective antagonist SCH-23390 (10 μM). Modulation experiments involved 5 min of baseline firing, either in the presence or absence of SCH-23390, followed by bath application of SKF-81297. Firing modulation was calculated by comparing the average number of action potentials evoked per stimulus in this baseline epoch with a 5-minute window starting 10 minutes after initial SKF-81297 application. All chemicals were purchased from Sigma or Tocris Bioscience.

### Two-photon microscopy

Two-photon imaging was performed on a custom microscope, as previously described (Chalifoux JR and AG Carter 2010). Briefly, a Ti:Sapphire laser (Coherent) tuned to 810 nm was used to excite Alexa Fluor 594 to image morphology with a 60x 1.0 NA objective (Olympus). Three-dimensional reconstructions of dendritic morphologies were performed using NeuronStudio (Wearne et al., 2005), while two-dimensional tracing of dendrites and axons for figures was performed using Neurolucida (MBF Bioscience). Dendrite analysis was performed by summing the total, apical or basal dendrite length contained within 10 μm concentric rings emanating from the soma and plotting these as a function of distance from the soma.

### Histology and fluorescence microscopy

Mice were anesthetized with a lethal dose of ketamine (25 mg/mL) and xylazine (0.25 mg/mL) and perfused intracardially with 0.01 M phosphate buffered saline (PBS) followed by 4 % paraformaldehyde (PFA) in 0.01 M PBS. Brains were fixed in 4 % PFA in 0.01 M PBS for 4-12 hours at 4°C. Slices were prepared at a thickness of 40-60 μm (Leica VT 1000S vibratome). For enhanced detection of tdTomato signal in D1-tdTomato mice slices were stained with antibodies against RFP. For antibody labeling, slices were washed once in PBS (0.01 M), once in PBS-T (0.2 % Triton-X100), then blocked in PBS-T with 1 % w/v bovine serum albumin (BSA) for one hour, all at room temperature. Primary antibody incubation (Rabbit anti-red fluorescent protein, 600-401-379, Rockland, 1:1000; Mouse anti-calretinin, MAB1568, Millipore, 1:1000; Mouse anti-parvalbumin, MAB1572, Millipore, 1:2000; Rat anti-somatostatin, MAB354, Millipore, 1:400) was performed at 4^°^C overnight. Slices were then washed 4x in PBS at RT before incubating with secondary antibody (Goat anti-rabbit Alexa 594, ab150080, AbCam, 1:400; Goat anti-rat Alexa 647, 21247, Fisher-Invitrogen, 1:200; Goat anti-mouse Alexa 647, ab150119, Abcam, 1:200) in PBS-T + BSA for 1 hour at room temperature. Slices were washed a further 3x in PBS before being mounted under glass coverslips on gelatin-coated slides using ProLong Gold antifade reagent with DAPI (Invitrogen). Whole-brain images were acquired using a slide-scanning microscope (Olympus VS120) with a 10x 0.25 NA or 20x 0.75 NA objective. Excitation wavelengths were 387, 485, 560 and 650 nm for DAPI, FITC, TRITC and Cy5, respectively. PFC images were acquired using a confocal microscope (Leica SP8) with 10x 0.4 NA or 20x 0.75 NA objective. Excitation wavelengths were 405, 488, 552 and 638 nm for DAPI, FITC, TRITC and Cy5, respectively. Image processing involved adjusting brightness and contrast using ImageJ (NIH). Cell counting was performed in a 400 x 1000 μm region of interest across the depth of the prelimbic prefrontal cortex.

### *In situ* hybridization

Mice were anesthetized with a lethal dose of ketamine (25 mg/mL) and xylazine (0.25 mg/mL) and perfused intracardially with chilled 0.01 M PBS. The brain was extracted and immediately submerged in isopentane cooled on dry ice. Tissue was coated in O.C.T. media (Tissue Tek) and stored in an airtight container at −80°C until sectioning. 10 μm sections were taken on a cryostat at −20°C and mounted on Superfrost Plus microscope slides (Fisher) and stored at −80°C. *In situ* hybridization of Mm-Drd1a-C2 and tdTomato-C3 probes was performed using a standard RNAscope protocol for flash frozen tissue from ACD bio. Slides were mounted under glass coverslips using ProLong Gold antifade reagent with DAPI (Invitrogen). Images were acquired using a confocal microscope (Leica SP8) with 20x 0.75 NA or 40x 1.3 NA oil immersion objective.

### Data analysis

Electrophysiology and imaging data were acquired using National Instruments boards and custom software written in MATLAB (MathWorks). Off-line analysis was performed using custom software written in Igor Pro (WaveMetrics). Input resistance was measured using the steady-state response to a −50 pA current injection for pyramidal neurons and −10 or −20 pA for interneurons. The membrane time constant (tau) was measured using exponential fits to these same hyperpolarizations. Voltage sag due to h-current was calculated by taking the minimum voltage in the first 200 ms, subtracting the average voltage over the final 100 ms, and dividing by the steady-state value. Spike frequency adaptation was calculated by calculating the ratio of the initial inter spike interval (ISI) and final ISI in response to a 500 ms depolarizing current pulse which evoked >5 action potentials. For cell counting as a function of layer, individual cells were assigned a distance from the midline (top of layer 1), binned in 25 μm increments across the depth of PFC, and then assigned into individual layers. Layers were defined based on peaks in neuron density (**Table 2**), which gave defined ranges for each layer (**Table 4**). Data was collected from at least 3 slices per animal, with a minimum of 3 mice per projection class / interneuron subtype. Cell-by cell mRNA puncta analysis was performed using a circular region of interest (17.5 μm in diameter) placed over the soma of individual neurons and manual counting of puncta for Drd1a and tdTomato.

### Experimental design and statistical analysis

Summary data are reported in the text and shown in figures as arithmetic mean ± SEM, unless otherwise stated. Statistical comparisons were performed in GraphPad Prism (version 7.0c) using a two-tailed non-parametric Mann-Whitney U test. Significance was defined as *p* < 0.05.

## RESULTS

### D1 receptors are expressed in excitatory and inhibitory cells in the PFC

We studied D1-receptor expressing (D1+) neurons using D1-tdTomato mice, in which expression of the red fluorescent protein tdTomato is driven by the D1 receptor promoter (**Fig. 1A**) (Ade KK *et al.* 2011). D1+ neurons were prominent in the prelimbic (PL) PFC, with most cells residing in layer 5 (L5) and 6 (L6) (**Fig. 1B & C**; L5 = 33 ± 3% of all D1+; L6 = 48 ± 3%, n = 8 mice). We also observed a small population of neurons located in superficial layers 1 (L1), 2 (L2) and 3 (L3) (**Fig. 1B & C**; L1 = 5 ± 1% of all D1+; L2 = 10 ± 1%; L3 = 4 ± 1%). Within each layer, D1+ neurons represented only a sub-population of cells, consistent with D1-R expression in a subset of pyramidal neurons (Seong HJ and AG Carter 2012).

**Figure 1:**
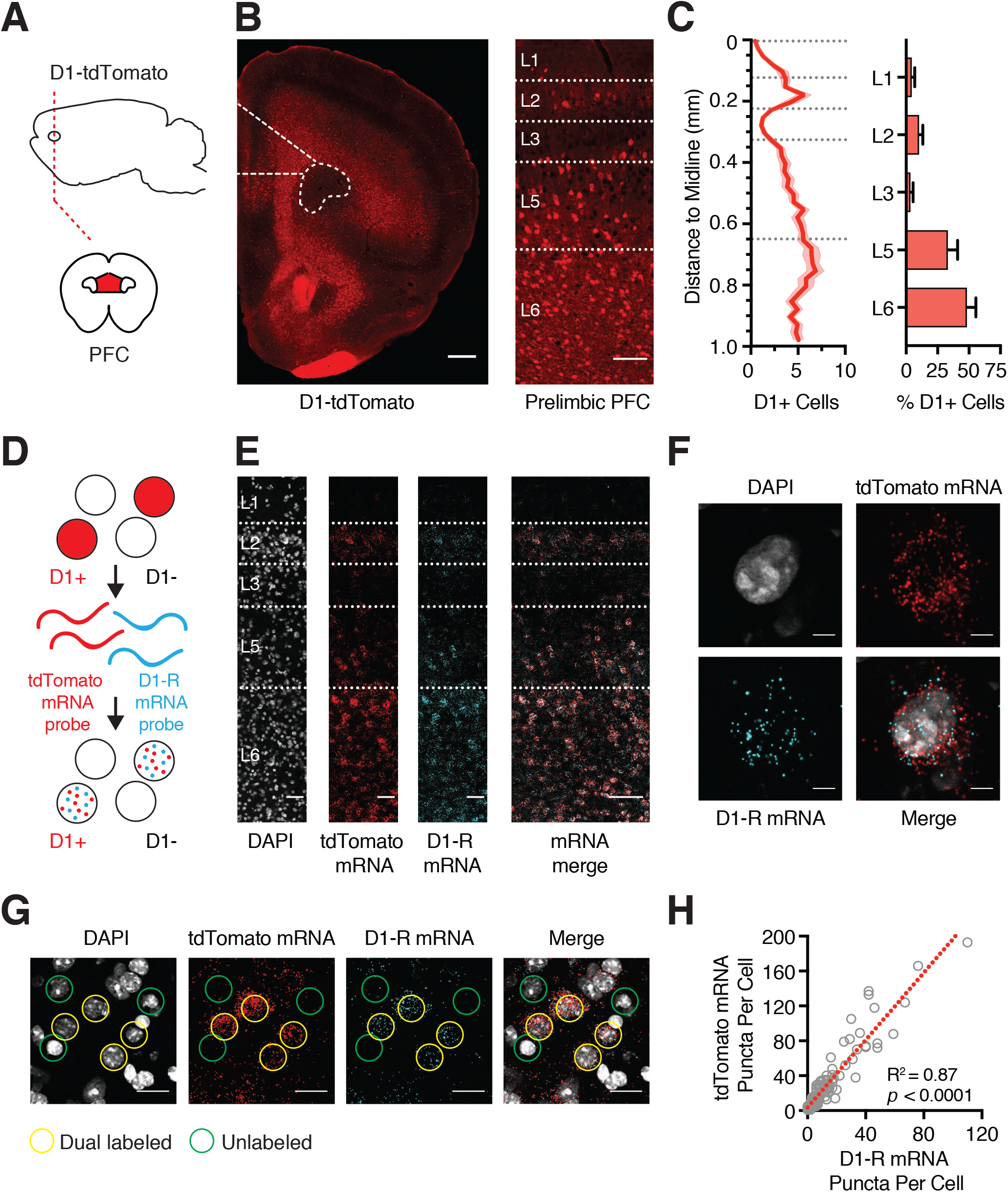
D1-tdTomato mice replicate endogenous D1-R expression. **A)** Schematic of coronal slice through the medial PFC (red). **B)** *Left*, Coronal slice showing D1-tdTomato expression in the PFC, highlighting the prelimbic subdivision in dotted outline. *Right*, Expanded view showing distribution of D1-tdTomato positive (D1+) neurons across layers of prelimbic PFC. Scale bars = 500 μm and 100 μm, respectively. **C)** Summary of D1+ neuron distribution as a function of distance from midline (left) and as a function of different layers (right). Bin size = 25 μm. Dashed lines represent laminar boundaries. **D)** Schematic of *in situ* hybridization process to distinguish tdTomato mRNA (red) and D1-R mRNA (blue) in single PFC neurons. **E)** *Left to right*, Distribution of DAPI-labeled cells (grey), tdTomato mRNA (red), D1-R mRNA (blue) and merged image, showing co-localization of mRNAs across layers of prelimbic PFC. Scale bars = 50 μm and 100 μm. **F)** Zoomed image showing co-localization of tdTomato and D1-R mRNA within an individual L5 PFC neuron. Scale bars = 5 μm. **G)** Co-localization of tdTomato and D1-R mRNA within a subset of deep layer PFC neurons. Examples of cells enriched for both mRNAs are highlighted in yellow, while poorly enriched cells are shown in green. Scale bars = 20 μm. **H)** Correlation of number of D1-R and tdTomato mRNA puncta within individual neurons. Dashed red line shows linear fit to the data. Values shown as mean ± SEM.

To validate that tdTomato faithfully replicates endogenous D1-R expression, we probed for D1-R (Drd1a) and tdTomato mRNAs using multiplex fluorescent *in situ* hybridization (**Fig. 1D**) (Wang F et al. 2012). The distributions of mRNA puncta for Drd1a and tdTomato were similar, and mirrored that observed for D1-tdTomato cells (**Fig. 1B & E**). On a cell-by-cell basis, there was strong overlap in expression levels of Drd1a and tdTomato mRNA (**Fig. 1F & G**), with strong correlation between the number of Drd1a and tdTomato positive puncta in individual neurons (**Fig. 1F**; n = 105 cells, from 3 mice, R^2^ = 0.87, *p* < 0.0001). These findings support the use of D1-tdTomato mice to explore D1+ neurons in the PFC.

Cortical neurons are broadly divided into either glutamatergic pyramidal neurons or GABAergic interneurons. To label glutamatergic neurons, we injected AAV-CaMKII-EGFP virus into the PFC of D1-tdTomato mice (**Fig. 2A**). We found that most D1+ neurons were also CaMKII+ (87 ± 2% of D1+ cells, n = 5 mice), indicating the majority are glutamatergic.

**Figure 2:**
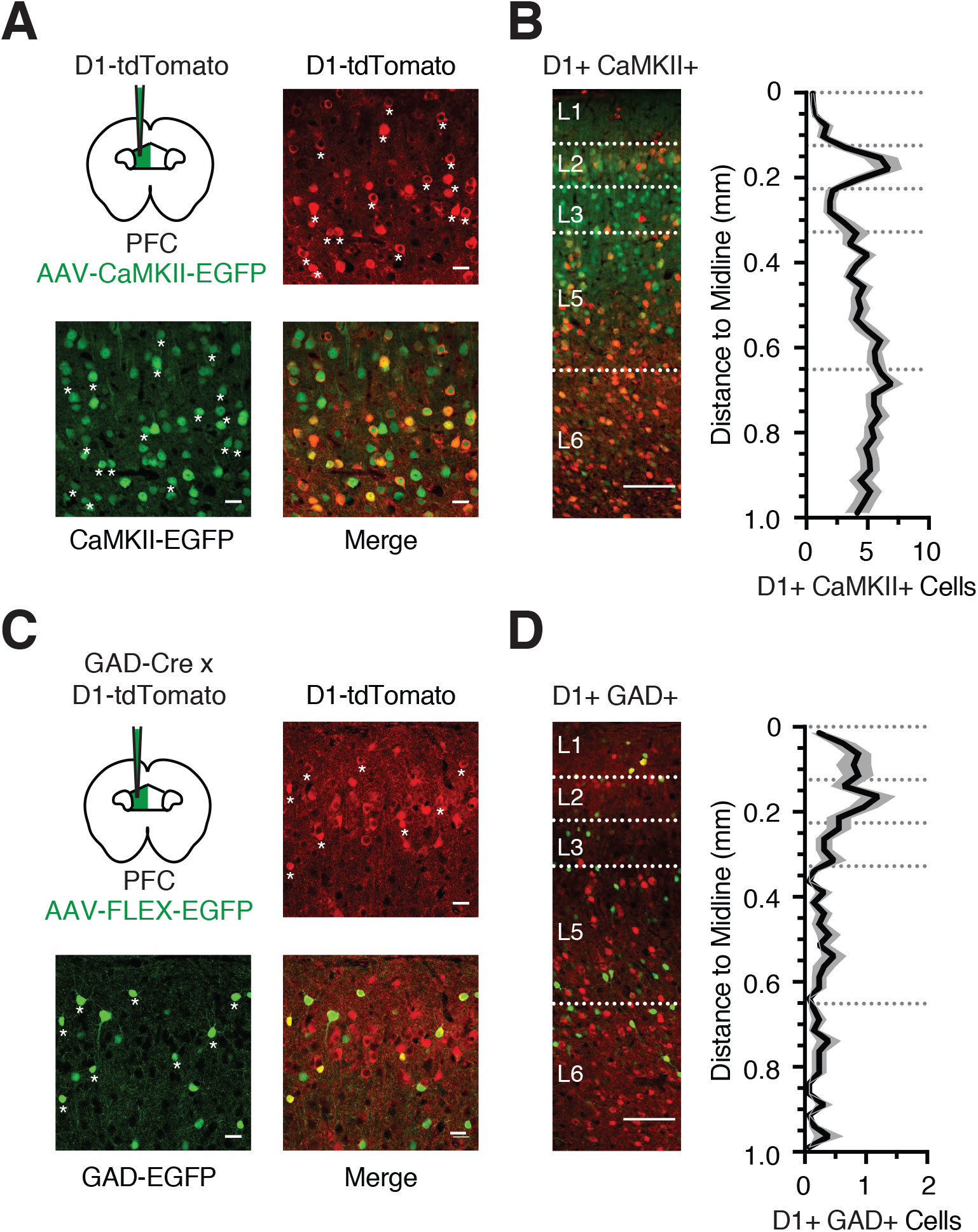
D1 receptors are expressed in pyramidal neurons and interneurons. **A)** Schematic for injection of AAV-CaMKII-EGFP into the prelimbic PFC of D1-tdTomato mouse, and confocal images from L5 showing D1-tdTomato expression (red), CaMKII-EGFP expression (green), and merged image. Asterisks indicate a subset of co-labeled neurons. Scale bar = 25 μm. **B)** *Left*, Distribution and overlap of D1+ (red) and CaMKII+ (green) neurons across layers of the prelimbic PFC. *Right*, Summary distribution of co-labeled D1+ CaMKII+ neurons as a function of distance from midline. Bin size = 25 μm. **C)** Similar to (A) for injection of AAV-FLEX-EGFP into GAD-Cre x D1-tdTomato mouse, showing D1-tdTomato expression (red), GAD-EGFP expression (green) and merged image from superficial layers. Asterisks indicate co-labeled neurons. Scale bar = 25 μm. **D)** *Left*, Distribution and overlap of D1+ (red) and GAD+ (green) neurons across layers of the prelimbic PFC. *Right*, Summary distribution of co-labeled D1+ GAD+ neurons of cell density as a function of distance from midline. Bin size = 25 μm. Values shown as mean ± SEM.

Accordingly, most co-labeled neurons were found in deep layers, similar to the overall D1+ population (**Fig. 2B**). Despite this strong overlap, we also observed some D1+ CaMKII-neurons, which could be cells that avoided viral transfection with the CaMKII virus, or alternatively a population of D1+ GABAergic interneurons that co-express GAD (glutamate decarboxylase). To test for the latter possibility, we crossed D1-tdTomato x GAD-Cre transgenic mice and injected Cre-dependent AAV-FLEX-EGFP virus into the PFC (**Fig. 2C**). Although most GAD+ neurons were D1-R negative (D1-), a population of co-labeled interneurons was present in superficial layers (**Fig. 2D**; 7.8 ± 0.1% of D1+ cells, n = 3 mice). Together, these findings indicate that the majority of D1+ CaMKII+ neurons are located in L5 and L6, whereas D1+ GAD+ interneurons are found in L1 and L2.

### D1+ pyramidal neurons have distinct properties and respond to D1-R activation

Pyramidal neurons in deep layers of cortex segregate into multiple subtypes based on their dendritic morphology and intrinsic physiology (Hattox AM and SB Nelson 2007; Dembrow NC *et al.* 2010; Thomson AM 2010; Anastasiades PG *et al.* 2018). We previously showed that D1+ and D1-pyramidal neurons in L5 of the juvenile PFC differ in their morphology and physiology (Seong HJ and AG Carter 2012). To extend these findings, we compared D1+ and D1-pyramidal neurons in both L5 and L6 of the adult PFC (**Fig. 3A & C**; D1+: L5 n = 7, L6 n = 8; D1-: L5 n = 8, L6 n = 7). We found sparse apical dendrites in L5 D1+ neurons (D1+ = 1827 ± 219 μm; D1-= 3448 ± 377 μm; *p* = 0.002) and L6 D1+ neurons (D1+ = 462 ± 323 μm; D1- = 2218 ± 336 μm; *p* = 0.01), with the latter often having multipolar or inverted dendrites (**Fig. 3A & C**). We also found different intrinsic properties in L5 D1+ neurons, which are more hyperpolarized, have higher input resistance, and minimal voltage sag (**Fig. 3B & Table 1**). Differences persisted for L6 D1+ neurons, which showed less voltage sag but similar resting potential and input resistance (**Fig. 3D & Table 1**). Interestingly, L6 D1-neurons also displayed minimal spike frequency adaptation, in contrast to the other cell types we recorded (**Fig. 3D & Table 1**).

**Figure 3:**
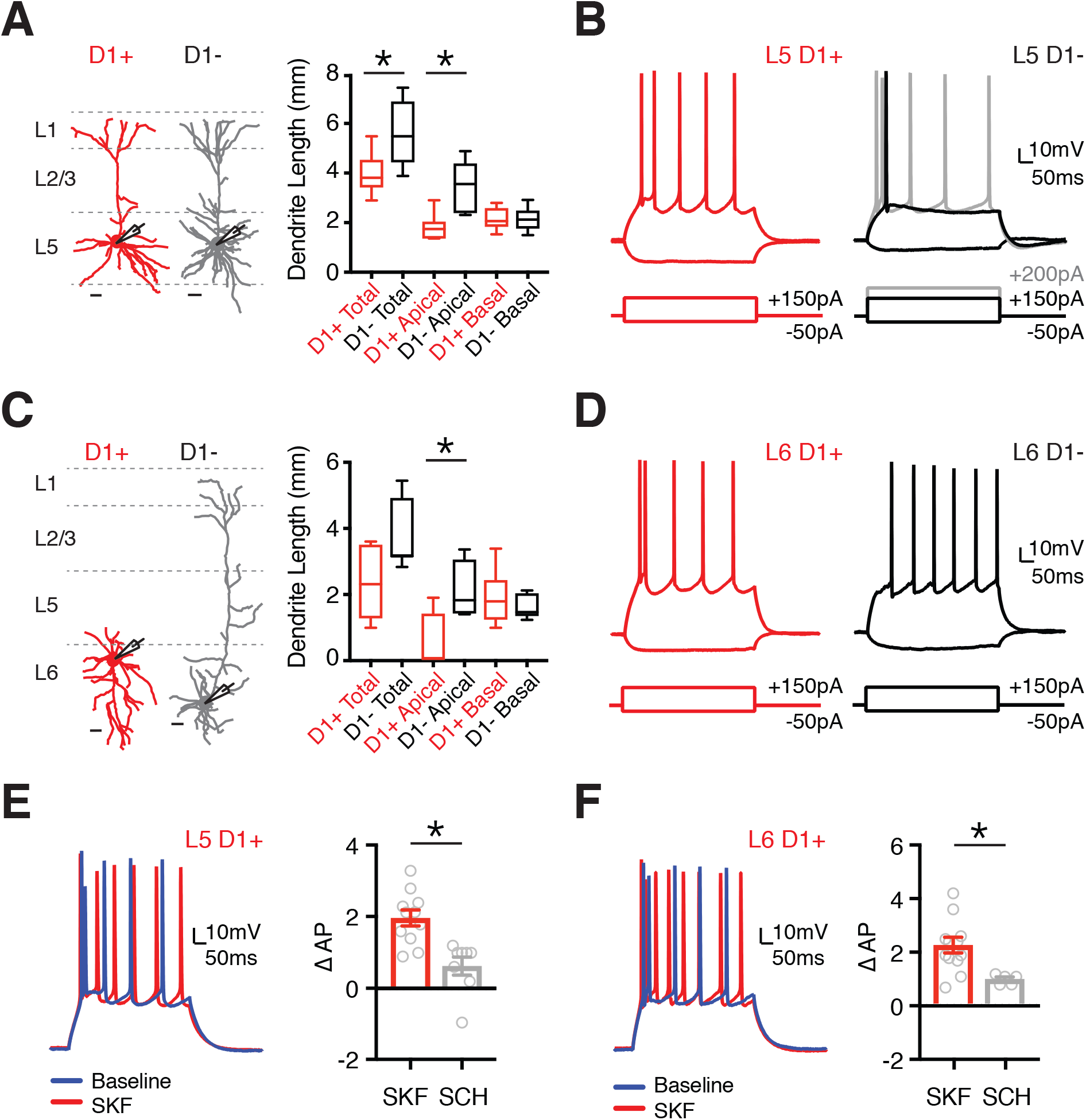
L5 and L6 D1+ pyramidal neurons are modulated by D1 receptors. **A)** *Left,* Dendrite reconstructions from 2-photon images of L5 D1+ (red) and L5 D1-(grey) pyramidal neurons. Scale bars = 50 μm. *Right*, Quantification of total, apical, and basal dendrite length for D1+ and D1-L5 pyramidal neurons. **B)** Intrinsic properties and AP firing of L5 D1+ (red) and L5 D1-(black / grey) pyramidal neurons in response to depolarizing and hyperpolarizing current steps. **C - D)** Similar to (A - B) for L6 D1+ (red) and L6 D1-(black) pyramidal neurons. **E)** *Left*, AP firing of L5 D1+ pyramidal neurons in response to depolarizing current step during baseline (blue) and after wash-in of the D1-R agonist SKF-81297 (10 μM) (red). *Right*, Summary of change in number of evoked APs (∆AP) recorded from L5 D1+ pyramidal neurons after application of either SKF or SCH + SKF. **F)** Similar to (E) for L6 D1+ pyramidal neurons. Values shown as median ± quartiles (A & C) or mean ± SEM (E & F). * p<0.05 *See also Table 1.*

**Table 1.**
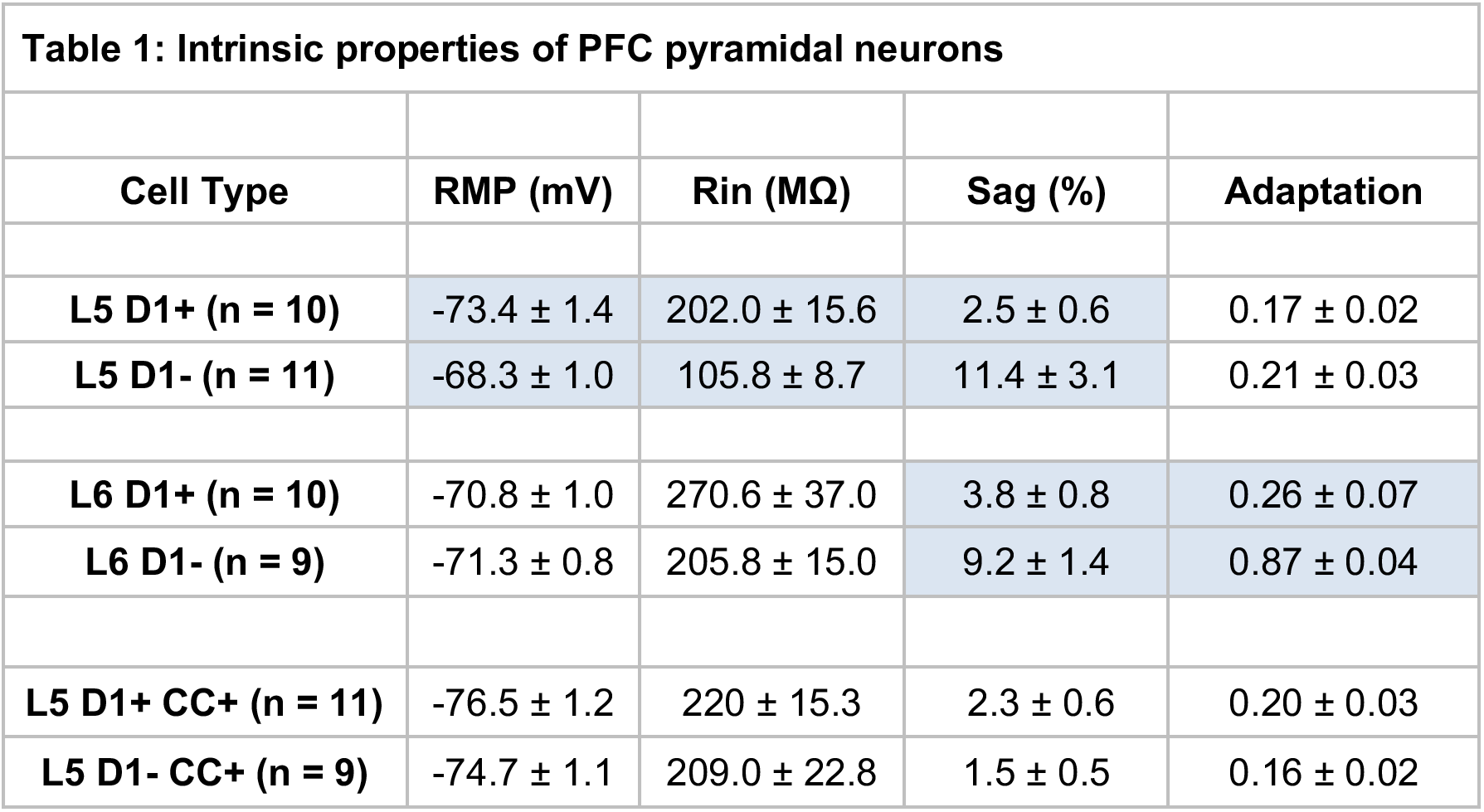
Resting membrane potential (RMP), input resistance (Rin), voltage sag due to h-current (Sag %) and adaptation ratio for L5 and L6 D1+ and D1-pyramidal neurons. Values in blue denote significance of p<0.05.

| Table 1: Intrinsic properties of PFC pyramidal neurons |  |  |  |  |
| --- | --- | --- | --- | --- |
| Cell Type | RMP (mV) | Rin (MΩ) | Sag (%) | Adaptation |
| L5 D1+ (n = 10) | -73.4 ± 1.4 | 202.0 ± 15.6 | 2.5 ± 0.6 | 0.17 ± 0.02 |
| L5 D1- (n = 11) | -68.3 ± 1.0 | 105.8 ± 8.7 | 11.4 ± 3.1 | 0.21 ± 0.03 |
| L6 D1+ (n = 10) | -70.8 ± 1.0 | 270.6 ± 37.0 | 3.8 ± 0.8 | 0.26 ± 0.07 |
| L6 D1- (n = 9) | -71.3 ± 0.8 | 205.8 ± 15.0 | 9.2 ± 1.4 | 0.87 ± 0.04 |
| L5 D1+ CC+ (n = 11) | -76.5 ± 1.2 | 220 ± 15.3 | 2.3 ± 0.6 | 0.20 ± 0.03 |
| L5 D1- CC+ (n = 9) | -74.7 ± 1.1 | 209.0 ± 22.8 | 1.5 ± 0.5 | 0.16 ± 0.02 |

Having explored the intrinsic properties of D1+ projection neurons, we assessed if they are modulated by D1-Rs. Dopamine receptors are known to regulate action potential (AP) firing in many brain regions, including the PFC (Gulledge AT and DB Jaffe 1998; Henze DA et al. 2000; Gorelova N et al. 2002; Seamans JK and CR Yang 2004). To rule out network effects, we studied modulation of AP firing in the presence of synaptic blockers. In L5 D1+ neurons, we found the D1-R agonist SKF-81297 (10 μM) increased firing, which was blocked by pre-incubation with the D1-R antagonist SCH-23390 (10 μM) (**Fig. 3E**; ∆AP: SKF = 2.0 ± 0.2, n = 11; SKF + SCH = 0.6 ± 0.3, n = 8; *p* = 0.0015). Similar regulation was seen for L6 D1+ neurons, with enhanced firing following SKF-81297, but not with SCH-23390 (**Fig. 3F**; ∆AP: SKF = 2.2 ± 0.3, n = 11; SKF + SCH = 0.9 ± 0.1, n = 5; *p* = 0.01). Together, these results indicate that L5 and L6 D1+ neurons are morphologically and physiologically distinct from adjacent D1-neurons, and that functional D1-Rs in these cells robustly enhance AP firing. Similar intrinsic physiology and morphology are observed for retrogradely labeled intra-telencephalic (IT) neurons (Hattox AM and SB Nelson 2007; Dembrow NC *et al.* 2010; Thomson AM 2010; Anastasiades PG *et al.* 2018), suggesting that projection neuron identity may help classify D1+ and D1-neurons in mouse PFC.

### Multiple classes of projection neurons contact long-range targets

To begin to explore projection neurons, we first determined the long-range targets of the PFC by injecting AAV-SynaptoTag, which labels axons in red and synapses in green at different target regions (Xu W and TC Sudhof 2013) (**Fig. 4A**; n = 3 mice). Monosynaptic target regions were distinguished from passing axons (red) by the co-localization of axonal and synaptic labeling (red and green). These regions included the contralateral PFC (cPFC), contralateral and ipsilateral claustrum (cCLA and iCLA), contralateral and ipsilateral striatum (cSTR and iSTR), mediodorsal (MD) and ventromedial (VM) thalamus, basolateral amygdala (BLA), and ventral tegmental area (VTA) (**Fig. 4A**). These various output pathways represent potential targets for D1+ projection neurons in the PFC.

**Figure 4:**
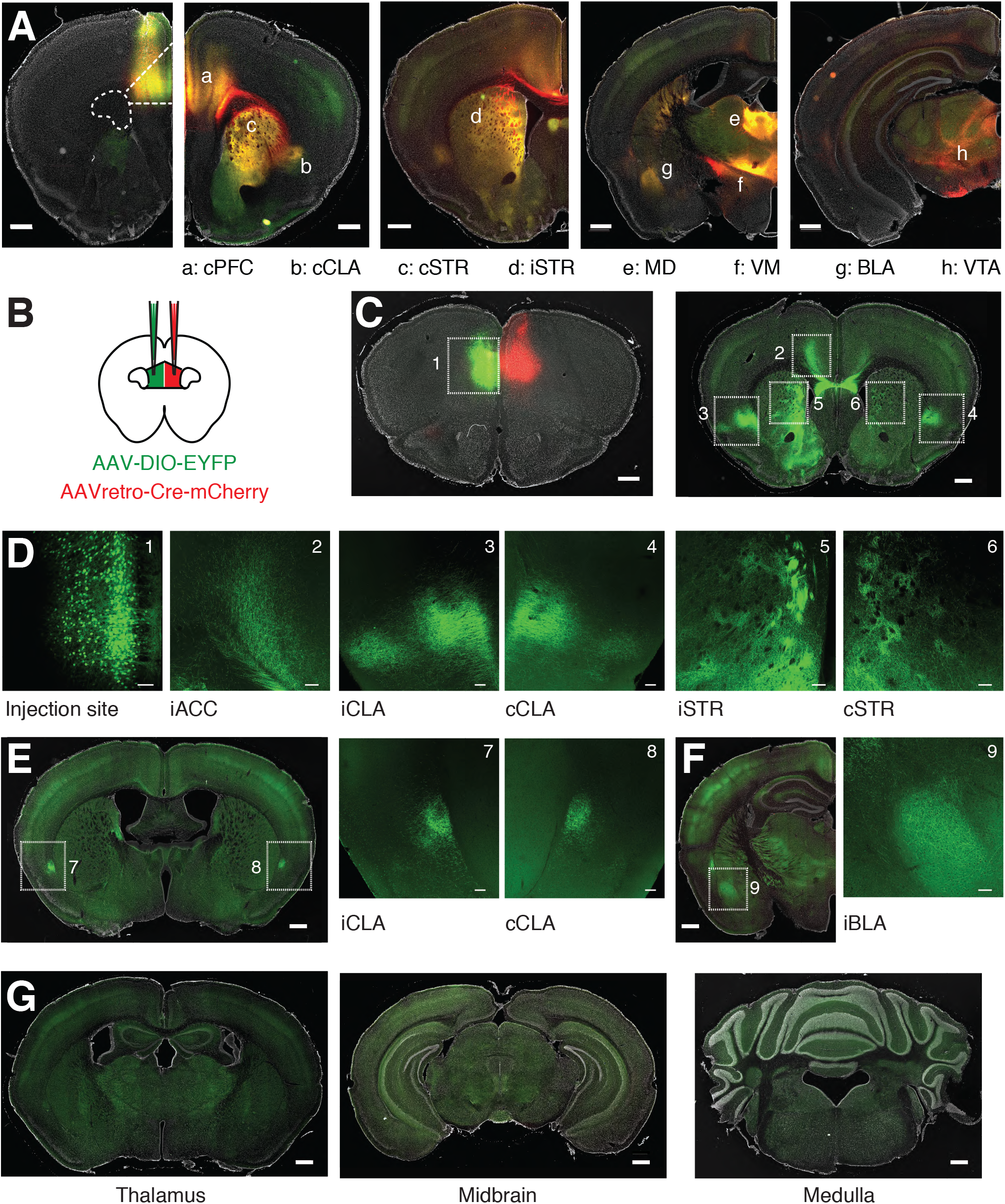
Characterization of PFC projection targets across the brain. **A)** *From left to right*, Representative injection of AAV-SynaptoTag, showing labeling of PFC axons (red), synaptic terminals (green) and their overlap (yellow) in (a) contralateral PFC (cPFC), (b) contralateral claustrum (cCLA), (c) contralateral dorsomedial striatum (cSTR), (d) ipsilateral dorsomedial striatum (iSTR), (e) mediodorsal thalamus (MD), (f) ventromedial thalamus (VM), (g) basolateral amygdala (BLA), and (h) ventral tegmental area (VTA). Scale bar = 500 μm. **B)** Schematic for injection of AAVretro-Cre-mCherry into the cPFC and AAV-DIO-EYFP into the ipsilateral (i)PFC. **C)** *Left*, Representative injection sites, showing EYFP-labeled CC neurons in iPFC, and location of AAVretro-Cre-mCherry in cPFC. *Right*, Distribution of CC neuron axons throughout multiple IT projection targets, including cortex, striatum and claustrum. Numbered boxes indicate zoomed imaging regions shown in (D). Scale bars = 500 μm. **D)** Confocal images of boxed regions in (C), highlighting labeled cell bodies and axons. *Left to right*, The AAV-DIO-EYFP injection site, iACC, iCLA, cCLA, iSTR, cSTR. Scale bars = 100 μm. **E)** *Left*, Distribution of CC neuron axons in caudal claustrum. Scale bar = 500 μm. *Right*, Confocal images of boxed regions highlighting labeled axons in both iCLA and cCLA. Scale bars = 100 μm. **F)** *Left*, Distribution of CC neuron axons in iBLA. Scale bar = 500 μm. *Right*, Confocal image of boxed region highlighting labeled axons in iBLA. Scale bar = 100 μm. **G)** *Left to right*, Absence of CC neuron axons from thalamus, midbrain and medulla. Scale bars = 500 μm.

Our physiology experiments indicate that D1+ neurons display IT properties, similar to cortico-cortical (CC) neurons projecting via the corpus callosum (Hattox AM and SB Nelson 2007; Dembrow NC *et al.* 2010; Thomson AM 2010; Anastasiades PG *et al.* 2018). However, pyramidal neurons can also send branching projections to multiple long-range targets (Wilson CJ 1987; Cowan RL and CJ Wilson 1994). To determine any additional targets of CC neurons, we next injected AAVretro-Cre-mCherry into the cPFC (Tervo DG et al. 2016), along with AAV-DIO-EYFP into the ipsilateral (i)PFC (**Fig. 4B & C**). EYFP+ CC neurons were distributed across layers (**Fig. 4D**), with labeled axons found in several distant brain regions, including the ipsilateral anterior cingulate cortex (iACC), iCLA, cCLA, iSTR and cSTR (**Fig. 4C & D**). CC neurons also projected to more caudal claustral regions (**Fig. 4E**) and sent axons to the iBLA (**Fig. 4F**), but not PT targets such as thalamus, midbrain and medulla (**Fig. 4G**). Interestingly, these findings are similar to recent reports of a genetically defined population of deep layer IT neurons (Nakayama H et al. 2018). Taken together, these data suggest that D1+ neurons may be IT cells that target the cortex, claustrum and striatum, motivating us to examine these projection neurons in more detail, including how these cell types segregate across different layers of the PFC.

### Projection neuron subtypes define distinct layers in PFC

Throughout cortex, subclasses of projection neurons display specific distributions across different layers (Gabbott PL *et al.* 2005; Oberlaender M et al. 2012; Oswald MJ et al. 2013; Harris KD and GM Shepherd 2015). To thoroughly determine these distributions in PFC, we independently injected Alexa-conjugated cholera toxin subunit B (CTB) into identified target regions (BLA n = 4, iSTR n = 3, cSTR n = 3, cPFC n = 3, cCLA n = 3, VTA n = 4, Pons n = 3, MD n = 4, VM n = 3). We observed retrogradely labeled projection neurons across layers 2-6 of PFC, with different classes distributed in a laminar-specific manner (**Fig. 5A**). Based on these bands of cell density, we were able to designate individual layers across the depth of PFC (**Fig. 5B & C**). Overall, we found that: L1 lacked projection neurons; L2 possessed a high density of cortico-amygdala (CA) neurons, but also a diverse array of other intra-telencephalic (IT) cells, including cortico-striatal (CS) neurons projecting to iSTR; L3 primarily comprised IT cells, with a greater density of CS neurons projecting to cSTR and cortico-cortical (CC) neurons. L5 could be divided into 3 distinct sublayers (Lorente de No R 1992): L5a contained a second band of CA neurons, the peak of CS neurons projecting to cSTR, and increased density of cortico-claustral (CCL) neurons; L5b possessed non-IT populations, including pyramidal tract (PT) and cortico-thalamic (CT) neurons, with PT neurons biased to upper L5b and less dense in lower L5b. Finally, L6 lacked PT neurons and contained both IT cells and a higher density of CT neurons. Together, these findings highlight the distribution and laminar structure of different projection neurons in the prelimbic PFC (**Fig. 5B & C**), extending on previous work in rat (Gabbott PL et al., 2005). Moreover, they allowed us to define specific boundaries for individual layers and sublayers in this agranular region of cortex (**Table 2**) and suggest that D1+ neurons located in L5 and L6 are likely to be CC or CCL cells.

**Figure 5:**
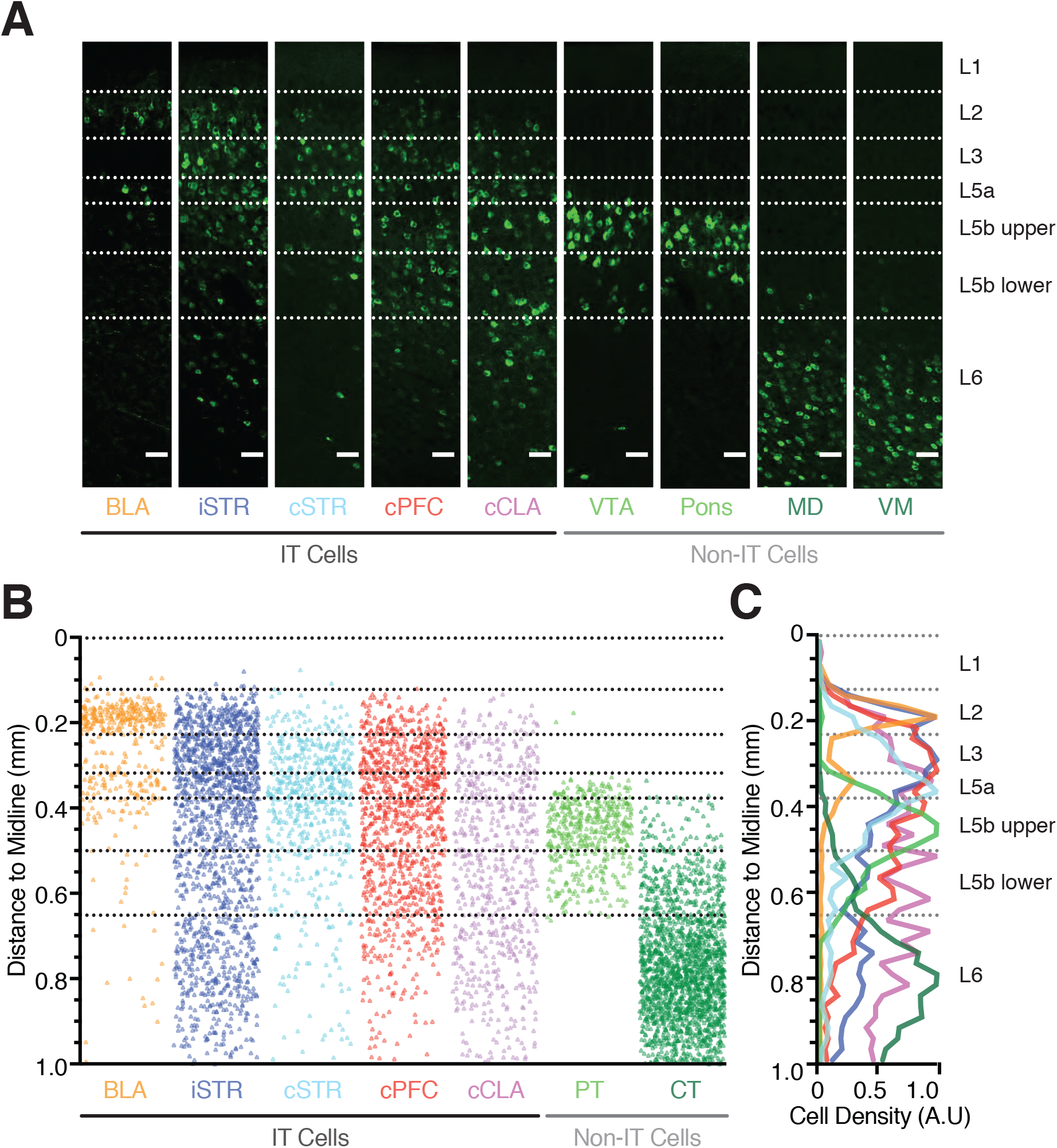
Laminar distributions of different projection neurons in the PFC. **A)** Confocal images showing the distribution of retrogradely labeled neurons across the depth of prelimbic PFC, following injection of fluorescently tagged Cholera Toxin subunit B (CTB) into BLA, iSTR, cSTR, cPFC, cCLA, VTA, pons, MD, or VM. Scale bars = 50 μm. **B)** Overlay of the position of all counted retrogradely labeled projection neurons as a function of distance to the midline. VTA and Pons are combined as pyramidal tract (PT) neurons, MD and VM are combined as cortico-thalamic (CT) neurons. **C)** Summary of peak-normalized cell density for projection neurons as a function of distance to the midline. *See also Table 2.*

**Table 2.** Designating individual layer boundaries based on the start and end of peak projection neuron density across multiple projection neuron populations.

| **Table 2: Projection neuron boundaries** |  |
| --- | --- |
| **L1 / L2 Border** | **Distance from Midline (μm)** |
| Start of BLA Cells | 125 ± 8 |
| **L2 / L3 Border** | **Distance from Midline (μm)** |
| End of 1<sup>st</sup> peak of BLA Cells | 219 ± 3 |
| End of 1<sup>st</sup> peak D1 Cells | 231 ± 5 |
| **L3 / L5a Border** | **Distance from Midline (μm)** |
| Start of 2<sup>nd</sup> peak of BLA Cells | 321 ± 9 |
| End of 2<sup>nd</sup> peak of BLA Cells | 376 ± 7 |
| **L5a / L5b upper Border** | **Distance from Midline (μm)** |
| Start of Pons Cells | 345 ± 3 |
| Start of VTA Cells | 375 ± 10 |
| **L5b upper / L5b lower Border** | **Distance from Midline (μm)** |
| End of peak cSTR Cells | 530 ± 8 |
| End of peak CT/PT dual labeled* | 492 ± 27 |
| **L5b lower / L6 Border** | **Distance from Midline (μm)** |
| End of VTA Cells | 664 ± 16 |
| End of Pons Cells | 627 ± 8 |

### D1 receptors are found in a subset of intra-telencephalic neurons

To determine which projection neuron populations express D1-R, we next examined co-localization of D1+ neurons and different populations of CTB-labeled (CTB+) neurons (**Fig. 6A**). For each of the retrogradely labeled populations (Fig. 4), we first quantified the percentage of D1+ neurons that are also CTB+ across all layers (% (D1+ CTB+) / D1+).

**Figure 6:**
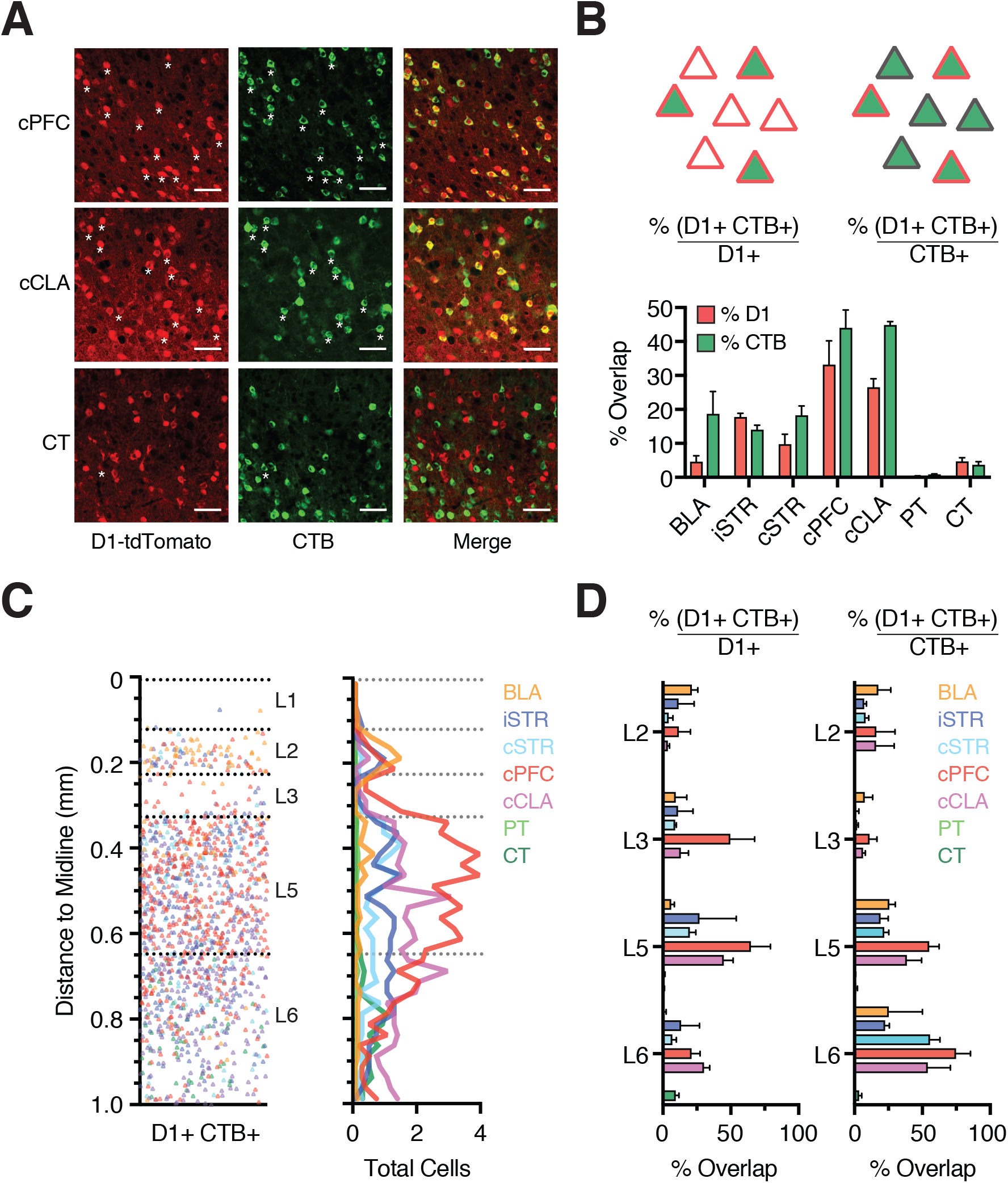
D1+ neurons in L5 and L6 are predominantly IT neurons. **A)** Confocal images showing overlap of D1+ neurons (red) and retrogradely labeled CTB+ neurons (green) that project to cPFC (top), cCLA (middle), or thalamus (CT, bottom). Scale bar = 50 μm. Asterisks indicate a subset of co-labeled neurons. Images taken from layer 5 (cPFC and cCLA) and layer 6 (CT). **B)** *Top left*, Schematic indicating the percentage of D1+ neurons across all layers that are retrogradely labeled (% (D1+ CTB+) / D1+). *Top right*, Schematic indicating the percentage of CTB+ neurons that are also D1+ neurons (% (D1+ CTB+) / CTB+). *Bottom*, Summary of both the percentage of D1+ neurons (% D1) and the percentage of CTB+ neurons (% CTB), respectively, for each of the projection classes shown in Fig. 5. **C)** *Left,* Overlay of all retrogradely labeled D1+ CTB+ co-labeled projection neurons as a function of distance to the midline. *Right,* Density plot of all D1+ CTB+ co-labeled projection neurons as a function of distance to the midline. Bin size = 25 μm. **D)** *Left,* Summary of percentage of D1+ neurons that are retrogradely labeled (% (D1+ CTB+) / D1+) as a function of layer. *Right,* Summary of percentage of CTB+ neurons that are also D1+ neurons (% (D1+ CTB+) / CTB+) as a function of layer. Values shown as mean ± SEM.

We found strong co-labeling for cells projecting to cPFC and cCLA (**Fig. 6A & B**; cPFC = 33 ± 7%, n = 3 mice; cCLA = 27 ± 4%, n = 3 mice), less co-labeling for cells projecting to BLA and striatum (**Fig. 6A & B**; BLA = 4.6 ± 1.8%, n = 4 mice; iSTR = 17.8 ± 1.1%, n = 3 mice, cSTR = 9.8 ± 2.9%, n = 3 mice), essentially no co-labeling for PT neurons projecting to pons or VTA, and a small yet consistently co-labeled population of L6 CT neurons projecting to MD and VM (**Fig. 6A & B**; PT = 0.4 ± 0.1%, VTA n = 4 mice, pons n = 3 mice; CT = 4.7 ± 1.1%, MD n = 4 mice, VM n = 3 mice) (Hoerder-Suabedissen A et al. 2018). Plotting (% (D1+ CTB+) / D1+) across the depth of PFC revealed projection-specific distributions (**Fig. 6C**), with CC and CCL neurons showing the strongest overlap across several layers, and particularly high co-labeling in deep layers (**Fig. 6D**).

Although this analysis identifies the main projection targets of D1+ neurons, it could miss cells that represent only a small proportion of the overall population, but themselves possess a high degree of D1-R enrichment. To account for this possibility, we also quantified the percentage of CTB+ neurons that are D1+ (% (D1+ CTB+) / CTB+). Approximately half of all neurons projecting to cPFC or cCLA were D1+ (**Fig. 6B**; cPFC = 44 ± 5%, cCLA = 45 ± 1%). However, this metric could be skewed by the fact that many of these neurons are located in L3, where D1+ neurons are largely absent. Accordingly, plotting (% (D1+ CTB+) / D1+) across layers revealed that a high percentage of CC and CCL neurons in L5 and L6 were D1+ (**Fig. 6D**; cPFC: L5 = 55 ± 7%, L6 = 75 ± 11%; cCLA: L5 = 39 ± 11%, L6 = 54 ± 17 %). Furthermore, around 20% of neurons that project to either BLA or striatum are also D1+ (BLA = 19 ± 7%, iSTR = 14 ± 1%, cSTR = 18 ± 3%), but very few neurons projecting via the pyramidal tract or to thalamus (PT = 0.9 ± 0.1%, CT = 3.7 ± 1%). These findings indicate that we did not overlook a key D1+ sub-type (**Fig. 6C & D**). Together, these results confirm that D1+ pyramidal cells are primarily IT neurons located in deep layers of the PFC, with prominent projections to cPFC and cCLA.

### D1 receptors modulate a sub-population of cortico-cortical neurons

Our data indicate that in L5 the majority of D1+ neurons are IT cells, while most D1-neurons are PT cells. However, despite significant overlap between D1-R expression and cPFC projecting cells, some CC neurons were observed to be D1-negative. This suggested modulation can be uncoupled from projection target, motivating us to compare D1+ and D1-CC neurons. Whole-cell physiology and dendritic reconstructions revealed similar morphology and physiology, with indistinguishable RMP, Rin, voltage sag and adaptation (**Fig. 7A-C & Table 1**; CC+ D1+ n = 11, CC+ D1-n = 9). However, while bath application of SKF enhanced the firing of CC+ D1+ neurons, it had no effect on CC+ D1-neurons (**Fig. 7D & E**; ∆AP: CC+ D1+ = 2.2 ± 0.4, n = 7; CC+ D1- = 0.5 ± 0.3, n = 4; *p* = 0.003). These results confirm there are at least two populations of CC neurons in L5, with only a subset modulated by D1-Rs. They also indicate that D1-R expression and projection target are not synonymous, such that not all CC neurons are D1+ neurons, and *vice versa*.

**Figure 7:**
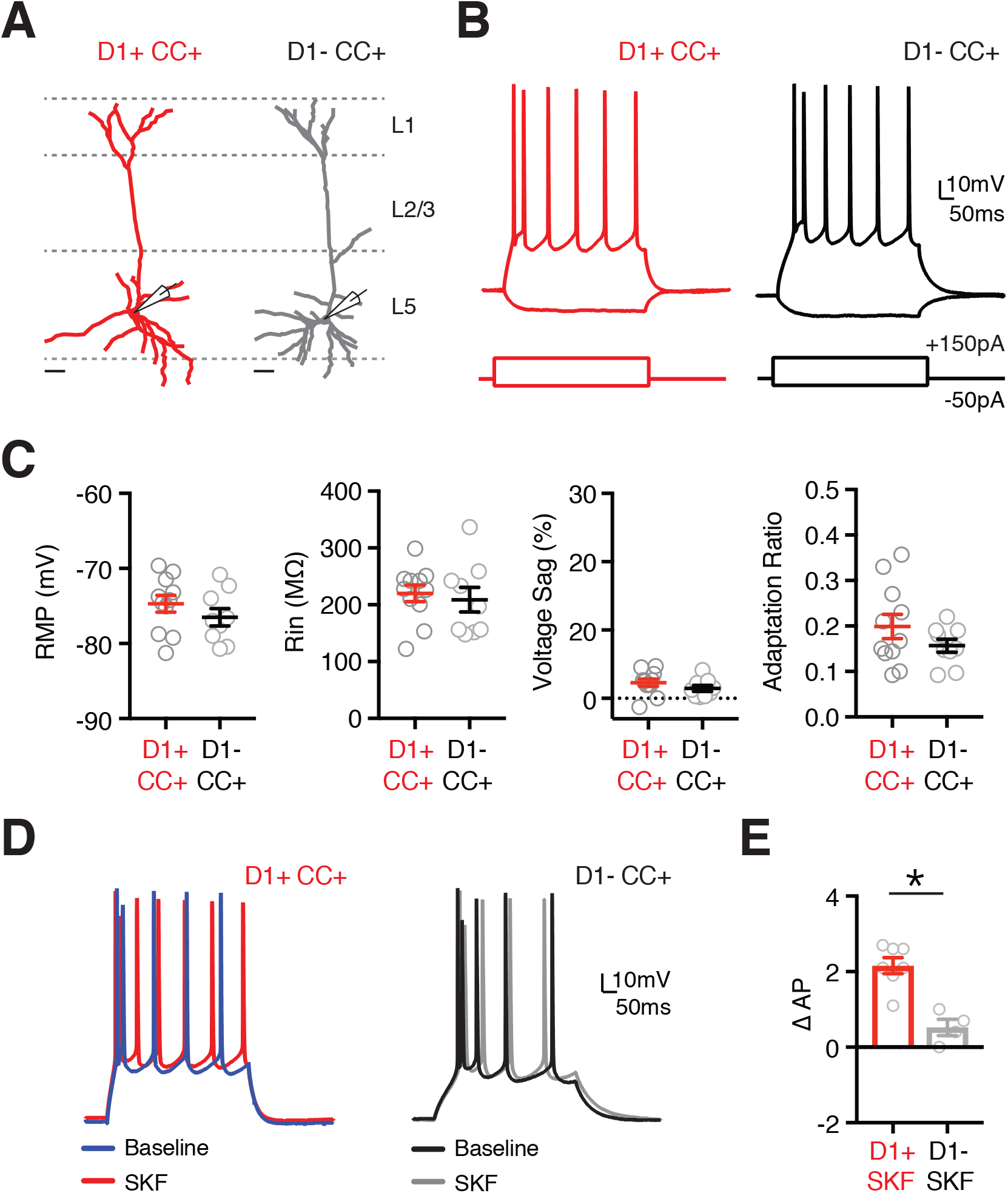
Heterogeneous modulation of CC neurons by D1 receptors. **A)** Dendrite reconstructions from 2-photon images of representative retrogradely labeled D1+ and D1-cortico-cortical (CC) neurons (D1+ CC+ in red, D1-CC+ in grey) whose cell bodies are located in layer 5 (L5). Scale bars = 50 μm. **B)** Intrinsic properties and AP firing of D1+ CC+ (red) and D1-CC+ (black) neurons, in response to depolarizing and hyperpolarizing current steps. **C)** Summary of resting membrane potential (RMP), input resistance (Rin), voltage sag (Sag %) and adaptation ratio in the two cell types. **D)** AP firing of D1+ CC (left) and D1-CC (right) neurons at baseline (blue / black) and in response to wash-in of the D1-R agonist SKF-81297 (10 μM) (red / grey). **E)** Summary of change in number of evoked APs (∆AP) recorded from D1+ CC+ and D1-CC+ neurons after application of SKF-81297 (10 μM). Values shown as mean ± SEM. * p<0.05 *See also Table 1.*

### D1 receptors are expressed in VIP+ interneurons

Our initial results indicated that, in addition to projection neurons, D1-Rs are expressed in a subset of GABAergic interneurons. Cortical interneurons are often segregated into distinct subtypes based on their physiology, morphology and expression of histochemical markers (Kubota Y and Y Kawaguchi 1994; Kawaguchi Y and Y Kubota 1996; Cauli B et al. 1997; Anastasiades PG *et al.* 2016). Three of these markers, parvalbumin (PV+), somatostatin (SOM+) and the serotonin receptor 3a (5HT3a+), label almost 100% of cortical GABAergic interneurons (Rudy B *et al.* 2011). The 5HT3a+ population is particularly diverse and contains a further major subset of interneurons that express vasoactive intestinal peptide (VIP+) (Lee S et al. 2010; Rudy B *et al.* 2011). PV+ and SOM+ interneurons primarily inhibit pyramidal neurons, while VIP+ neurons inhibit other interneurons and are engaged in disinhibitory networks. To determine D1-R overlap within these populations, we first crossed PV-, SOM-, 5HT3a-, and VIP-Cre lines with the D1-tdTomato line to produce interneuron specific double transgenic mice. To selectively label cortical interneuron subtypes, we then injected AAV-FLEX-EGFP into the PFC of each mouse line (**Fig. 8A**). Consistent with studies from other regions of cortex (Gonchar Y *et al.* 2007; Lee S *et al.* 2010), we found that PV+ and SOM+ interneurons were distributed across layers 2-6, whereas 5HT3a+ and VIP+ interneurons were largely biased to superficial layers (**Fig. 8A & B**). This anatomical characterization provides insight into the laminar distributions of the main GABAergic interneuron populations in the mouse PFC. Given that overlap of D1+ and GAD+ is most prominent in superficial layers, these results suggest that D1-Rs are selectively expressed in VIP+ and/or 5HT3a+ interneurons.

**Figure 8:**
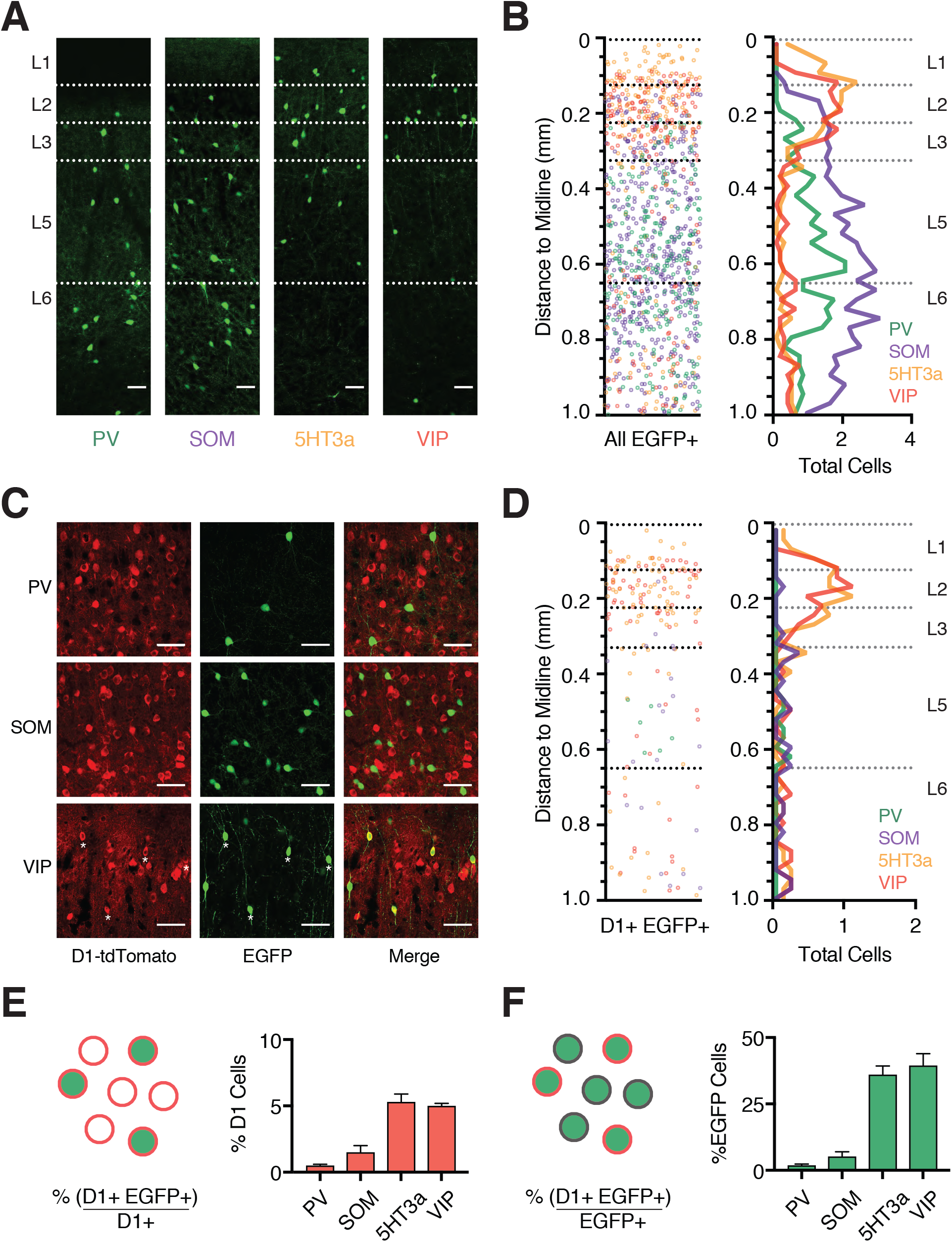
D1 receptors are expressed in a sub-population of interneurons. **A)** Confocal images of EGFP expression in the prelimbic PFC of PV-, SOM-, 5HT3a-and VIP-Cre x D1-tdTomato double transgenic mouse lines injected with AAV-FLEX-EGFP virus. Scale bar = 100 μm. **B)** *Left,* Overlay of the position of all EGFP+ interneurons as a function of distance to the midline. *Right,* Summary showing the average distribution of labeled PV+, SOM+, 5HT3a+ and VIP+ interneurons as a function of distance from the midline. Bin size = 25 μm. **C)** Confocal images showing D1+ (red), EGFP+ (green), and merged image, showing overlap in VIP+ interneurons (bottom) but not PV+ interneurons (top) or SOM+ interneurons (middle). Scale bar = 50 μm. Asterisks indicate co-labeled neurons. Images taken from layer 5 (PV+ and SOM+) and layer 2 (VIP+). **D)** *Left,* Overlay of the position of all D1+ EGFP+ co-labeled interneurons as a function of distance to the midline. *Right,* Summary showing the average distribution of D1+ EGFP+ co-labeled interneurons as a function of distance from the midline. Bin size = 25 μm. **E)** *Left,* Schematic highlighting the percentage of D1+ neurons which are co-labeled with EGFP (% (D1+ EGFP+) / D1+). *Right*, Summary as a function of interneuron subtype. **F)** Similar to (E), but for percentage of EGFP+ interneurons that are also D1+ (% (D1+ EGFP+ / EGFP+). Values shown as mean ± SEM.

We next examined the co-labeling of D1+ neurons within each interneuron population (**Fig. 8C & D**), using a similar analysis to that described for projection neurons above. Surprisingly, we observed minimal co-labeling of D1+ neurons and either PV+ or SOM+ interneurons (**Fig. 8C-E**; (% (D1+ EGFP+) / D1+): PV+ = 0.5 ± 0.1%, n = 3 mice; SOM+ = 1.5 ± 0.5%, n = 3 mice). In contrast, we observed substantial co-labeling of D1+ neurons and both 5HT3a+ and VIP+ interneurons (**Fig. 8C-E**; (% (D1+ EGFP+) / D1+): 5HT3a+ = 5.3 ± 0.6%, n = 3 mice; VIP+ = 5.0 ± 0.2%, n = 3 mice). Plotting the distribution of dual labeled cells as a function of distance to midline revealed pronounced 5HT3a+ and VIP+ co-labeling in superficial layers (**Fig. 8D**). Furthermore, the percentage of D1+ VIP+ interneurons was similar to that of D1+ GAD+ cells in our initial pan-interneuron experiments (Fig. 2). Given that the VIP+ population is a major subset of 5HT3a+ interneurons (Rudy et al., 2011), these findings suggest VIP+ interneurons comprise the majority of D1+ interneurons in the PFC. Interestingly, quantifying the number of EGFP+ interneurons that are also D1+ showed that less than half of 5HT3a+ and VIP+ interneurons are D1+ (**Fig. 8F**; (% (D1+ EGFP+) / EGFP+): 5HT3a+ = 36.1 ± 3.2%, VIP+ = 39.5 ± 4.4%), while also confirming minimal co-labeling of PV+ and SOM+ interneurons (**Fig. 8F**; (% (D1+ EGFP+) / EGFP+): PV+ = 1.9 ± 0.5%, SOM+ = 5.2 ± 1.8%). This was not an artifact of using Cre lines, as similar results were obtained with antibodies against PV and SOM (% (D1+ PV+) / PV+ = 4.1 ± 0.6%, n = 3 mice; % (D1+ SOM+) / SOM+ = 5.9 ± 1.0%, n = 3 mice). Together, these findings indicate that D1-Rs are expressed in a sub-population of VIP+ interneurons, which are primarily located in superficial layers of PFC.

### D1 receptors modulate a sub-population of VIP+ interneurons

VIP+ interneurons are highly diverse, comprising multiple distinct morphological and electrophysiological subtypes (Kawaguchi Y and Y Kubota 1996; Miyoshi G et al. 2010; Pronneke A et al. 2015; He M et al. 2016). To further characterize D1+ VIP+ interneurons, we next injected AAV-FLEX-EGFP into D1-tdTomato x VIP-Cre double transgenic mice, and performed targeted current-clamp recordings from cells in superficial layers (**Fig. 9A**). When considering all VIP+ interneurons, we observed many of the previously described firing patterns, including irregular-spiking (IS), non-fast spiking (NFS) and fast-adapting (fAD) subtypes (**Fig. 9A & Table 3**) (Miyoshi G *et al.* 2010). Interestingly, IS neurons were exclusively contained within the D1+ VIP+ population, forming a substantial proportion of D1+ VIP+ but not D1-VIP+ cells (**Fig. 9B**; D1+ VIP+: n = 9/14, D1-VIP+: n = 0/9). VIP+ interneurons that have IS firing properties typically have bipolar morphologies, co-express calretinin (CR), and target other interneurons to mediate disinhibition (Acsady L et al. 1996; Lee S et al. 2013; He M *et al.* 2016). Consistent with these findings, D1+ VIP+ interneurons had bipolar morphologies (**Fig. 9A**) and were frequently co-labeled with calretinin (**Fig. 9C**; n = 3). The proportion of D1+ VIP+ interneurons that were CR+ (67 ± 6 %) was very similar to the proportion of D1+ VIP+ interneurons with IS firing properties (64% of total). These findings indicate that D1-Rs are particularly enriched in a specific sub-population of VIP+ interneurons, which mediate disinhibition across cortex.

**Figure 9:**
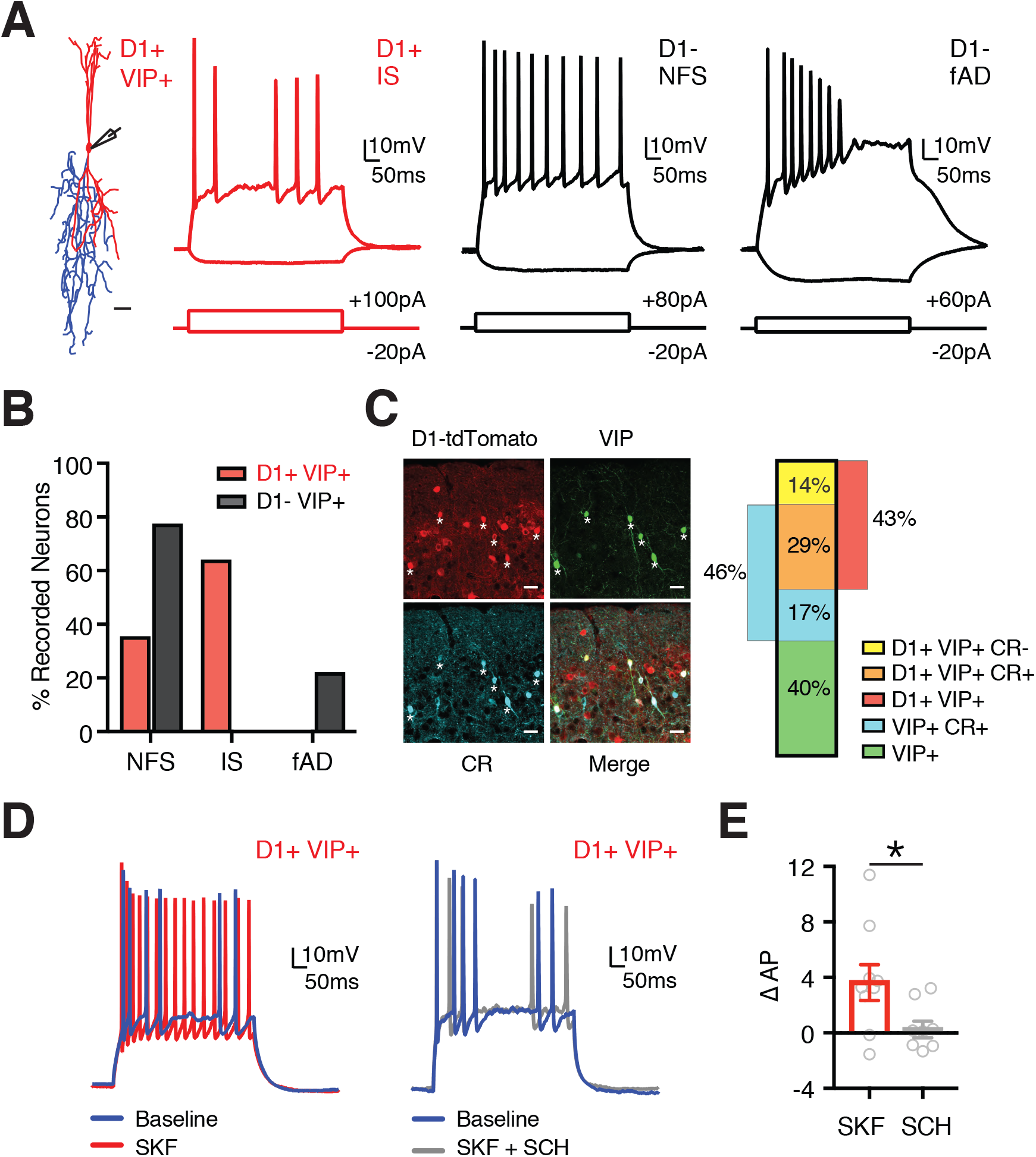
A specific subclass of D1+ VIP+ interneurons is modulated by D1-Rs. **A)** *Left*, Representative morphology of a D1+ VIP+ interneuron, with soma and dendrites in red and axon in blue. Scale bar = 25 μm. *Right*, Summary of responses to depolarizing and hyperpolarizing current steps for D1+ VIP+ and D1-VIP+ interneurons in superficial layers of prelimbic PFC. Intrinsic properties showing D1+ irregular spiking (IS), D1-non-fast spiking (NFS) and D1-fast-adapting (fAD) firing patterns. **B)** Summary of percentage of D1+ VIP+ (red) and D1-VIP+ (black) interneurons that exhibit NFS, IS, or fAD firing patterns. **C)** *Left,* Confocal images from superficial layers of D1+ (red), VIP+ (green), calretinin antibody staining (CR, cyan), and merged image. Scale bar = 25 μm. Asterisks indicate triple-labeled neurons. *Right,* Summary of percentages of labeled cells. **D)** Firing properties of D1+ VIP+ interneurons in baseline (blue) and following wash-in of either SKF alone (red, left) or SKF with SCH (grey, right). **E)** Summary of change in AP firing in D1+ VIP+ interneurons following wash-in of either SKF alone (red) or SKF with SCH (grey). Values shown as mean ± SEM. * p<0.05 *See also Table 3.*

**Table 3.** Resting membrane potential (RMP), input resistance (Rin), voltage sag due to h-current (Sag), membrane time constant (Tau), action potential (AP) height, maximum firing frequency (Max Freq.), afterhyperpolarization (AHP) amplitude, and AHP time to peak, for individual VIP+ interneuron populations in superficial layers of PFC.

| **Table 3: Intrinsic properties of PFC VIP+ interneurons** |  |  |  |  |  |  |  |  |
| --- | --- | --- | --- | --- | --- | --- | --- | --- |
| **Cell Type** | **RMP (mV)** | **Rin (MΩ)** | **Sag (%)** | **Tau (ms)** | **AP Height (mV)** | **Max Freq. (Hz)** | **AHP (mV)** | **AHP (ms)** |
| **IS (n = 9)** | -65.5<br>± 2.3 | 650.9<br>± 72.3 | 1.5<br>± 1.3 | 26.9<br>± 3.1 | 64.3<br>± 2.4 | 28.6<br>± 4.3 | 8.8<br>± 1.6 | 3.5<br>± 0.1 |
| **NFS (n = 12)** | -68.2<br>± 1.3 | 620.8<br>± 48.3 | 0.8<br>± 1.0 | 27.2<br>± 3.1 | 63.0<br>± 2.7 | 32.2<br>± 2.3 | 8.0<br>± 1.5 | 4.3<br>± 0.2 |
| **fAD (n = 2)** | -69.1 | 702.6 | 2.7 | 29.3 | 58.6 | 13 | 3.0 | 4.5 |

Finally, we examined D1-R modulation of D1+ VIP+ interneurons, in the presence of synaptic blockers to prevent network activity. We found that wash-in of SKF-81297 (10 μM) alone strongly enhanced the firing of these cells, whereas wash-in of SKF-81297 in the presence of SCH-23390 (10 μM) had no effect on firing (**Fig. 9D & E**; ∆AP: SKF = 3.8 ± 1.4, n = 9, SKF + SCH = 0.4 ± 0.6, n = 8; *p* = 0.04). Together, these findings indicate that D1-Rs also strongly enhance the firing properties of D1+ VIP+ interneurons.

## DISCUSSION

We have determined the cell-and layer-specific expression of D1 dopamine receptors in the mouse PFC. We found that D1-Rs robustly modulate subsets of pyramidal neurons and GABAergic interneurons. We showed significant overlap between deep layer D1+ neurons and cells that project throughout the telencephalon. Additionally, we observed a sub-population of superficial D1+ VIP+ GABAergic interneurons, which are known to inhibit other interneurons. Together, our results highlight the specificity with which D1-Rs exert influence on both excitatory and disinhibitory micro-circuits in the mouse PFC.

Our data indicate that most D1+ neurons reside in L5 and L6, which parallels the increased density of both dopaminergic axon terminals and dopamine receptors in deeper layers of PFC (Berger B et al. 1976; Santana N *et al.* 2009; Van De Werd HJ et al. 2010). Most D1+ neurons are also glutamatergic, consistent with ultrastructural observations, which indicate the presence of D1-Rs at glutamatergic presynaptic terminals and postsynaptic spines (Smiley JF et al. 1994; Paspalas CD and PS Goldman-Rakic 2005), where they may also function to regulate synaptic responses (Gao WJ et al. 2001; Urban NN et al. 2002). Moreover, L5 and L6 D1+ neurons are modulated by D1-Rs, leading to enhanced AP firing, consistent with our previous work in juvenile mice (Seong HJ and AG Carter 2012). However, while D1-Rs have been proposed to enhance persistent firing of L3 pyramidal neurons in primates (Paspalas CD et al. 2013), they are conspicuously absent from L3 in mouse. This may represent differences in cortical architecture across species, with L3 functioning as the primary thalamo-recipient layer in mouse (Collins et al., 2018). Alternatively, the enhancement in firing in superficial layers may be mediated by alternative mechanisms, for example VIP mediated disinhibition.

Defining the laminar boundaries in the PFC and other frontal cortices has been challenging, because landmarks like L4 are absent (Uylings HB et al. 2003). Our anatomical results revealed the distribution of numerous cell types in the PFC, extending on work in rats (Gabbott PL *et al.* 2005). Superficial layers contain most of the 5HT3a+ and VIP+ interneurons, which also populate superficial layers in sensory cortices (Gonchar Y *et al.* 2007; Lee S *et al.* 2010). They are also enriched in cortico-amygdala (CA) neurons (Gabbott PL *et al.* 2005; Little JP and AG Carter 2013) and contain a high density of IT neurons (Wilson CJ 1987; Otsuka T and Y Kawaguchi 2011; Oswald MJ *et al.* 2013; Anastasiades PG *et al.* 2018). Deep layers of PL can be delineated by the increased density of subcortical projection neurons (Molnar Z and AF Cheung 2006; Oswald MJ *et al.* 2013; Harris KD and GM Shepherd 2015). Interestingly, L5 can be subdivided into 3 distinct sublayers based on the relative density and identity of PT neurons (Lorente de No R 1992; Gabbott PL *et al.* 2005; Collins DP et al. 2018). This analysis elaborates on recent studies (DeNardo LA et al. 2015; Clarkson RL *et al.* 2017), and provides clarity on the extent of L3 and sublayers of L5. Together, these findings highlight the complexity of the mouse PFC, detailing its laminar structure (**Tables 2 & 4**).

**Table 4.** Designating lower layer boundaries after averaging across projection neuron populations shown in Table 2. Analysis range indicates distances from midline used to divide data into individual layers using 25 μm bins. WM = white matter.

| **Table 4: Layer boundaries** |  |  |
| --- | --- | --- |
|  | **Lower Bound**<br>**($\mu\text{m}$ from Midline)** | **Analysis Range**<br>**($\mu\text{m}$ from Midline)** |
| **L1** | $125 \pm 8$ | 0 – 125 |
| **L2** | $227 \pm 3$ | 126 – 225 |
| **L3** | $321 \pm 9$ | 226 – 325 |
| **L5a** | $376 \pm 7$ | 326 – 375 |
| **L5b upper** | $511 \pm 5$ | 376 – 500 |
| **L5b lower** | $648 \pm 7$ | 501 – 650 |
| **L6** | WM | 651 – 1000 |

Using retrograde tracing, we identified L5 and L6 D1+ neurons as IT cells, which project to both ipsilateral and contralateral cortex (Mercer A et al. 2005; Brown SP and S Hestrin 2009; Morishima M et al. 2011). Interestingly, we also observed a high percentage of deep layer D1+ neurons projecting laterally, towards the claustrum. Consistent with D1-R expression in cortico-claustral (CCL) neurons, L6 D1+ neurons have inverted pyramidal morphologies (Bueno-Lopez JL et al. 1991; Mendizabal-Zubiaga JL et al. 2007). Projections from frontal cortex to claustrum are more prevalent than those from sensory cortices (Atlan G et al. 2017; Brown SP et al. 2017) and may play an important role in cognitive functions (Smith JB and KD Alloway 2014; White MG et al. 2018). Given that both dopamine and the claustrum are implicated in hallucinations and attention deficits (Goll Y et al. 2015; Brown SP *et al.* 2017), the expression of D1-Rs in CCL neurons may be significant for schizophrenia and ADHD (Goll Y *et al.* 2015; Grace AA 2016). In addition to D1+ CC and CCL neurons, we observed some D1+ CA neurons, which regulate feeding (Land BB *et al.* 2014). Finally, we observed a subset of D1+ CS neurons, consistent with IT neurons sending collaterals to iSTR and cSTR (Wilson CJ 1987; Cowan RL and CJ Wilson 1994). Together with our AAVretro-Cre axon tracing experiments, these findings suggest that D1-Rs impact a distributed IT network projecting to multiple targets, potentially leading to diverse effects on cognition and behavior.

Our results indicate multiple populations of IT neurons exist within PFC, with distinct dopamine receptor expression profiles. The highest density of IT neurons is in L3, the only layer where D1-Rs were largely absent. Within L5, not all IT cells express D1-Rs, with L5 D1-CC neurons insensitive to D1-R agonists. One possibility is that these cells may be D3+ neurons, whose firing is suppressed by D3 receptors (Clarkson RL *et al.* 2017). Alternatively, D1-R expression may fluctuate within the IT cell-class, either over the course of development, in response to activity, or salient stimuli (Brenhouse HC et al. 2008; Zhao Y et al. 2017). In the future, it will be interesting to assess the functional significance of individual IT neuron sub-classes within PFC, and determine the importance of dopamine modulation on these diverse populations of projection neurons (Otsuka T and Y Kawaguchi 2011; Hirai Y et al. 2012; Ueta Y et al. 2013; Yamashita T et al. 2013). Moreover, PT neurons are another class of D1-projection neurons whose physiology is influenced by D2 receptors (Gee S *et al.* 2012). These findings highlight the importance of both cell-type and layer in interpreting the impact of neuromodulators, and may account for some of the heterogeneity that was initially observed for dopamine modulation in unlabeled neurons (Penit-Soria J et al. 1987; Geijo-Barrientos E and C Pastore 1995; Gulledge AT and DB Jaffe 1998; Zhou FM and JJ Hablitz 1999; Gulledge AT and DB Jaffe 2001; Seamans JK and CR Yang 2004). They may also explain some of the varied effects of dopamine receptor agonists and antagonists on PFC-dependent behaviors (Floresco SB and O Magyar 2006; St Onge JR et al. 2011; Jenni NL *et al.* 2017).

In addition to pyramidal neurons, we observed that superficial layers also contain a population of D1+ GABAergic interneurons. Previous reports from primate PFC suggest that PV+ interneurons may express D1-Rs (Muly EC, 3rd *et al.* 1998;Glausier JR *et al.* 2009), but the degree of co-labeling reported varies widely (Le Moine C and P Gaspar 1998; Paspalas CD and PS Goldman-Rakic 2005; Tritsch NX and BL Sabatini 2012). Surprisingly, we observed minimal co-labeling between D1 receptors and either PV+ or SOM+ interneurons, suggesting these inhibitory circuits are not directly modulated by D1-Rs. One possibility is that there are pronounced differences in D1-R expression between species. Another interesting possibility is that PV+ interneurons may instead express the D5 receptor, as observed in the striatum (Centonze D et al. 2003; Oda S *et al.* 2010; Tritsch NX and BL Sabatini 2012).

Instead, D1-Rs are primarily found in superficial VIP+ interneurons, which show irregular spiking firing properties, co-express calretinin, and are strongly modulated by D1-Rs, leading to enhanced AP firing. Throughout cortex, VIP+ interneurons mediate disinhibition by inhibiting SOM+ interneurons in the local circuit (Pfeffer CK et al. 2013; Pi HJ et al. 2013; Karnani MM et al. 2016). In the PFC, recent studies highlight an important role for VIP+ activity in working memory tasks (Kamigaki T and Y Dan 2017), consistent with previous modeling studies (Wang XJ *et al.* 2004). Because dopamine levels increase during cognitive tasks (Watanabe M et al. 1997; Phillips AG et al. 2004), our findings provide a mechanism linking elevated PFC dopamine with VIP+ interneuron activity through D1-Rs. Moreover, engaging disinhibitory circuits could explain why superficial networks are enhanced by D1-Rs, even though receptors are not prominent in pyramidal cells. Previous studies indicate that VIP+ interneurons are also under the control of other neuromodulators, including serotonin, acetylcholine and noradrenaline (Beaulieu C and P Somogyi 1991; Smiley JF and PS Goldman-Rakic 1996; Paspalas CD and GC Papadopoulos 1999; Lee S *et al.* 2010; Letzkus JJ *et al.* 2011; Rudy B *et al.* 2011). Our findings provide further support for the idea that VIP+ mediated disinhibition may be a common circuit mechanism utilized by many neuromodulatory systems (Wester JC and CJ McBain 2014).

Together, our data provide a detailed overview of D1-R expression in both excitatory and inhibitory neurons of the mouse PFC. By increasing the firing of L5 D1+ IT cells projecting across the corpus callosum, D1-R activation may increase communication between hemispheres, which plays an important role in delay period activity (Li N et al. 2016). Activating these neurons may also regulate activity within the local network, where CC neurons make contacts onto other CC neurons, as well as CT and PT cells (Mercer A *et al.* 2005; Brown SP and S Hestrin 2009). Moreover, by increasing the activity of VIP+ interneurons, D1-Rs can disinhibit the local network, which is known to play an important role in cortical function, including within the PFC (Wang XJ *et al.* 2004; Garcia Del Molino LC et al. 2017; Kamigaki T and Y Dan 2017). Thus, D1-Rs can activate both excitatory circuits in deep layers and disinhibitory circuits in superficial layers. These findings help explain the role of D1-Rs in the PFC, with important implications for understanding dopamine modulation in cognitive processing and related neuropsychiatric disorders.

## ACKNOWLEDGEMENTS

We thank members of the Carter lab for helpful discussions and comments on the manuscript. We thank Susan Sheng, Mian Hou and Claudia Farb for help with immunocytochemistry and histology. This work was supported by NIH R01 MH085974 (AGC). The authors declare no financial conflicts of interest.

